# A Fast Adaptive Algorithm for Computing Whole-Genome Homology Maps

**DOI:** 10.1101/259986

**Authors:** Chirag Jain, Sergey Koren, Alexander Dilthey, Adam M. Phillippy, Srinivas Aluru

## Abstract

**Motivation:** Whole-genome alignment is an important problem in genomics for comparing different species, mapping draft assemblies to reference genomes, and identifying repeats. However, for large plant and animal genomes, this task remains compute and memory intensive.

**Results:** We introduce an approximate algorithm for computing local alignment boundaries between long DNA sequences. Given a minimum alignment length and an identity threshold, our algorithm computes the desired alignment boundaries and identity estimates using kmer-based statistics, and maintains sufficient probabilistic guarantees on the output sensitivity. Further, to prioritize higher scoring alignment intervals, we develop a plane-sweep based filtering technique which is theoretically optimal and practically efficient. Implementation of these ideas resulted in a fast and accurate assembly-to-genome and genome-to-genome mapper. As a result, we were able to map an error-corrected whole-genome NA12878 human assembly to the hg38 human reference genome in about one minute total execution time and < 4 GB memory using 8 CPU threads, achieving significant performance improvement over competing methods. Recall accuracy of computed alignment boundaries was consistently found to be > 97% on multiple datasets. Finally, we performed a sensitive self-alignment of the human genome to compute all duplications of length ≥ 1 Kbp and ≥ 90% identity. The reported output achieves good recall and covers 5% more bases than the current UCSC genome browser's segmental duplication annotation.

**Availability:** https://github.com/marbl/MashMap.

## 1 Introduction

Algorithms for inferring homology between DNA sequences have undergone continuous advances for more than three decades, mainly in the direction of achieving better accuracy to compare distant genomes, as well as better compute efficiency to scale with growing data. Up until the last decade, reconstruction of a complete reference genome through sequencing and assembly was deemed a major landmark in genomics (Lander *et al.*, 2001; Venter *et al.*, 2001). However, it did not take long for high-throughput sequencing technologies to fuel populationwide genomics projects through low-cost genome assemblies (e.g., the Genome 10K project, Haussler *et al.*, 2009). Analysis of these new genome assemblies, for both population-scale biological studies and timely diagnosis in clinical settings, requires faster and memory-efficient algorithms for facilitating whole-genome comparisons.

It is well-known that computing local alignments using an exact dynamic programming algorithm at the whole-genome scale is computationally prohibitive. This bottleneck motivated the development of seed-and-extend based genome aligners. Within the seed-and-extend paradigm, the two common approaches adopted to compute exact matches are either implemented using a hash table for *k*-mers (e.g., Altschul et *al*., 1997; Ma *et al.*, 2002; Schwartz *et al.*, 2003) or suffix trees and its variants (Delcher *et al.*, 1999; Brudno *et al.*, 2003; Bray *et al.*, 2003; Vyverman *et al.*, 2013; Marçais G *et al.*, 2018). A third category includes cross-correlation based algorithms (e.g., Satsuma by Grabherr *et al.*, 2010). However, these approaches still remain computationally intensive. For instance, Nucmer (Kurtz *et al.*, 2004) and LAST (Kielbasa *et al.*, 2011), two widely used genome-to-genome aligners, require 10 or more CPU hours to align a human genome assembly to a human reference genome.

The primary motivation behind this work is to develop a new genome-to-genome mapping algorithm that is faster and memory-efficient while maintaining accuracy on par with sensitive aligners. We seek a problem formulation that also provides a convenient handle for users to specify how diverged the input genomes are, based on their knowledge of which organisms are being compared, expected quality of genome assembly, and sensitivity requirements of any further downstream biological analysis.

The inspiration behind our algorithmic strategy stems from recent developments in techniques for long-read analyses. MinHash-based estimation of Jaccard similarity of *k*-mer sets between DNA sequences has been adopted for state-of-the-art long read genome assembly (Koren *et al.*, 2017), whole-genome distance computation (Ondov *et al.*, 2016; Jain *et al.*, 2017b), and long read mapping (Jain *et al.*, 2017a). Through our previous work Mashmap (Jain *et al.*, 2017a), we demonstrated that a MinHash-based approximate mapping algorithm can compute long-read mapping boundaries with accuracy on par with alignment-based methods, while exhibiting two orders of magnitude speedup. Mashmap operates by assuming an error-distribution model, links alignment identity to Jaccard similarity, and provides sufficient probabilistic guarantees on output sensitivity. However, this algorithm is limited to end-to-end mapping of input sequences, which makes it impractical for contig mapping or *split-read* mapping. Here, we introduce new algorithmic strategies to compute local alignment boundaries for both whole-genome and split-read mapping applications.

Given minimum identity and length requirements for local alignments, we formulate the characteristics of output we intend to compute. Our new algorithm internally makes use of our previous end-to-end approximate read mapping framework (Jain *et al.*, 2017a) by applying it to nonoverlapping substrings of the query sequence. We mathematically show that all valid local alignment boundaries, which satisfy the user-specified alignment identity and length thresholds are reported, with high probability. Further, we formulate a heuristic to prioritize mappings with higher scores. We leverage the classic plane-sweep technique from computational geometry to develop an *O(n* log *n)* algorithm to solve the filtering problem, with n being the count of total mappings.

We demonstrate the practical utility of our algorithm Mashmap2 by evaluating accuracy and computational performance using real data instances, which include mapping human genome assemblies and ultralong reads to the human reference genome, and sensitive self-alignment analysis of the human genome. We compared the performance of Mashmap2 against a recent fast alignment-free method Minimap2 (Li, 2017) and the widely used alignment-based method Nucmer (Kurtz *et al*., 2004). Mashmap2 operates in about a minute and 4 GB memory, including both indexing and mapping stages, to map human genome assembly to a reference when given minimum alignment identity and length requirements of 95% and 10 Kbp respectively. This makes it the most resource-efficient software for genome-to-genome mapping, especially with respect to the memory-usage. This performance is achieved while maintaining output sensitivity percentage in the high 90s. We also demonstrate its ability to compute all ≥ 1 Kbp long segmental duplications in the human genome with high accuracy. We expect the performance and sensitivity guarantees provided by our algorithm will allow fast evaluation of draft assemblies versus a reference genome, scalable construction of whole-genome homology maps, and rapid split-read mapping of long reads to large reference databases.

## 2 The Mashmap2 Algorithm

We designed Mashmap2 to enable fast computation of homology maps between two sequences or a sequence and itself. It consists of two algorithmic components. The first computes approximate boundaries and alignment scores for all pairs of substrings that exceed a user specified length and identity threshold. The second applies a novel filtering algorithm to optionally weed out redundant, paralogous mappings.

### 2.1 Computing Local Alignment Boundaries

Consider all local mappings of the form *Q[i..j]* between sequences *Q* (query) and *R* (reference) of length l_0_ or more, such that *Q[i..j]* aligns with a substring of R with per-base error-rate ≤ *ϵ_max_* and |*j — i* + 1| ≥ *l_0_*. Alignment algorithms have quadratic time complexity, therefore an exact evaluation of the local mappings between all possible substring combinations will require *O* ( | *Q* |^3^*R* |^3^ ) time. As such, solving this problem exactly is computationally prohibitive for typical sizes of real datasets. Instead of explicitly computing all such structures, we seek at least one seed mapping of length *l_0_/2* along the path of each optimal alignment. Doing so, while maintaining high sensitivity and sufficient specificity will allow computation of the local alignments efficiently using an appropriate alignment algorithm.

In our approach, we leverage our previous alignment-free end-to-end read mapping algorithm, designed for mapping noisy long reads (*Jain et al., 2017a*). This allows us to benefit from its attractive properties including probabilistic guarantees on quality, and algorithmic and space efficiency. We continue to assume the same error model that was used in this work, also restated here. We assume that alignment errors, i.e, substitutions and indels in a valid alignment occur independently and follow a Poisson distribution. We also simplify by assuming that *k*-mers are independent entities in sequences. For a given per-base error rate threshold *ϵ_max_*, the read-mapping algorithm reports all target mapping coordinates and identity estimates of a read in the reference, where it aligns end-to-end with *≤ ϵ_max_* per-base error rate, with high probability. This is achieved by linking Jaccard coefficient between the fc-mer spectra of the read and its mapping region to the alignment-error rate, under the assumed error distribution model.

#### Proposed Algorithm

We first split the query sequence *Q* into *l_0_/2* sized non-overlapping fragments. If a substring of *Q*, say *Q_sub_*, of length ≥ *l_0_* aligns against a substring of *R* with *ϵ ≤ ϵ_max_* per-base error rate, then there is at least one *l_0_/2* sized fragment that maps end-to-end along the optimal alignment path with *h ϵ · l_0_/2* expected errors. Accordingly, the read mapping routine in Mashmap can be used to map each fragment with *ϵ_max_* error-rate cutoff. Note that at least 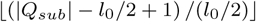 ≥ 1 query fragments completely span *Q_sub_* (Figure 1). Let *p* be the probability that a fragment is mapped to the desired target position on the reference, compinluted as described by Jain *et al.* (2017a). Probability of reporting at least one seed mapping along the optimal alignment is given by 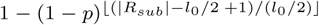. We show that these probability scores are sufficiently high, between 0.92 and 1.00 for alignment error rate thresholds *ϵ_max_* 10% and 20% respectively (Figure 2).

**Fig. 1.**
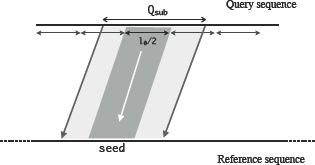
A local alignment depicting the inclusion of a length *lo√2* fragment of the query sequence.

**Fig. 2.**
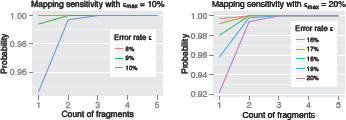
Probability of mapping at least one seed fragment for two different error-rate thresholds *ϵ_max_* = 10%, 20%. As true error rate e decreases, the probability values accordingly improve as expected. Similarly, longer alignments spanning more fragments are more likely to be reported. Most importantly, all the sensitivity scores are consistently above 90%. To compute the probability values, sketch size for Minhash based Jaccard estimation was assumed as 200, and the *k*-mer size was set to 16. These parameters are internally set by Mashmap (Jain et al., 2017a).

The above seed matches and their alignment identity estimates are further processed to compute approximate local boundaries and their scores. After computing all seed matches, matches which involve consecutive query sequence fragments are merged together if they are mapped closely in the same order on the reference sequence. Suppose mappings from the consecutive query fragments *q_i_,q_i_+_1_,…,qj* are mapped to reference positions with begin positions *p_0_,p_1_,…,pj-i* respectively, then they are grouped together as a local alignment segment if *p_0_ < p_1_ ≤… ≤ *Pj—i*, and*pk+_1_* — *pk < l_0_*, [0 < k < j —* i]. The alignment boundaries are estimated as the first and last mapping offsets of the group. The corresponding alignment scores are estimated as their average identity estimate multiplied by the sum of the fragment lengths. We use these alignment boundaries and the scores as input to a subsequent filtering algorithm.

### 2.2 A Geometric Algorithm for Filtering Alignments

Large mammalian genomes and plant genomes have abundant repetitive sequences. As a consequence, a large fraction of inferior mappings are reported due to paralogous genomic segments or false positive mappings resulting from simple sequence repeats. Furthermore, from a biological perspective, closely examining all alternative mappings may not be feasible. Therefore, different strategies are adopted to identify biologically relevant outputs. We formulate a filtering heuristic for our mapping application, and develop an optimal *O(n* log *n)* algorithm to solve it. We also prove that Ω(*n* log *n*) runtime is necessary to solve this problem. The effectiveness of this algorithm on real genomic data is demonstrated later, in the Results section.

#### 2.2.1 Problem Formulation

Suppose all output mappings of a query sequence are laid out as weighted segment intervals, with the alignment scores used as weights (Figure 3). We propose the following filtering heuristic: a segment is termed redundant if and only if it is subsumed by higher scoring segments at all of its positions. In practice, there can be multiple alignments with equal scores. Therefore, segment scores are allowed to be non-unique.

A sub-optimal *O*(*n*^2^) algorithm for solving the above problem can be readily developed by doing an all to all comparison among the segments. However, it would lead to practically slow implementation for typical input sizes. The formulated filtering problem bears resemblance to the line segment intersection test problem for which Shamos and Hoey (1976) gave a classic *O*(*n* log *n*) algorithm using plane-sweep technique. Accordingly, we summarize their algorithm next, and subsequently describe the modifications made to solve the above filtering problem.

**Fig. 3.**
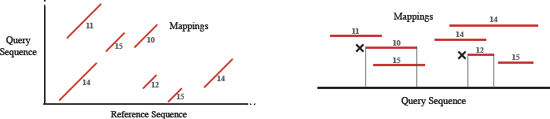
Left figure is a toy example to illustrate line segments corresponding to multiple local alignments obtained between query and reference sequence, similar to a dot-plot. Each alignment segment is labeled with an alignment score. Now, suppose we wish to filter best mappings for the query sequence. These segments can be considered as weighted intervals over the query sequence (right figure). In the above case, two intervals marked with a cross are completely subsumed by higher scoring intervals, and therefore, will be labeled as redundant by our filtering heuristic.

#### 2.2.2 The Shamos-Hoey Algorithm

Similar to the filtering problem, the problem of detecting whether n segments have an intersecting pair has a trivial *O*(*n*^2^) solution. Shamos and Hoey solved this problem using a plane-sweep based *O*(*n* log *n*) algorithm. The algorithm defines an ordering between segments in the 2D plane. The main loop of the algorithm conceptually *sweeps* a vertical line from left to right, and while doing so, the sweep-line status data-structure L dynamically holds segments which intersect the sweep-line.

The sweep-line halts at 2*n* endpoints of the input segments, and the order of segments in 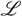 is evaluated to detect any intersection. For efficiency, this algorithm chooses a balanced tree to implement the sweep-line status 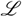. As such, it spends O(log *n*) time at each halting point, and therefore, the total runtime is bounded by *O(n* log *n*). This algorithm is popular not only for its theoretical and practical efficiency, but also for ease of implementation.

In our problem as well, evaluating segments which intersect the vertical sweep-line at 2*n* endpoints is sufficient to identify all *good* (nonredundant) segments. However, evaluating all intersecting segments at each endpoint is inefficient, and again leads to a quadratic algorithm. Therefore, we devise a new ordering scheme among segments which will enable us to evaluate only a subset of intersecting segments at each endpoint to bound the runtime.

#### 2.2.3 Proposed Algorithm for Alignment Filtering

The Shamos-Hoey algorithm requires several modifications for solving the filtering problem. We define an order between segments as following: Between two segments, the segment with higher score is considered as greater, but if the scores are equal, then the segment with the larger starting position is considered as greater. This particular ordering helps avoid redundant computations, and will be crucial for bounding the runtime later.

Similar to the Shamos-Hoey algrithm, we also use a height-balanced Binary Search Tree (BST) as the data-structure for the sweep-line status 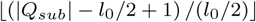, which tracks the segments that intersect the vertical sweep line. 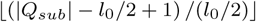 is required to support the following operations in our algorithm:

1. *insert(s).* Insert segment *s* into 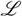.
2. *delete(s).* Delete segment s from 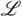.
3. *mark_good*(). Mark all segments with highest score as good in 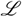.

Note that the *insert* and *delete* operations are naturally supported in O(logn) time in BSTs, whereas the *mark_good* function can be realized as a sequence of *maximum* and *predecessor* operations. If there are k segments with equal and highest scores in 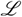, the function *mark_good* uses *O(k* log *n*) time. With the data-structures and the operations defined above, we give an outline of the complete filtering procedure in Algorithm 1. The main loop of the algorithm iterates over the 2*n* segment endpoints, which is analogous to the sweep line moving from left to right, halting at the 2*n* points. In each iteration, we update the sweep-line status 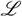 so that it holds the segments which intersect the sweep line, and mark the highest-scoring segments as *good* using the *mark_good* function.

##### Lemma 1

*Algorithm 1 solves the filtering problem correctly.*

Proof. Consider a function *S* : N → {0,1}^N^ from positions in the query sequence to subsets of segments {1, 2,…, *N*}. A segment s_i_ *ϵ {S(pos)}* if and only if it is among the highest scoring segments which overlap with the query sequence at position *pos.* Clearly, a union of all subsets in the domain of function S equals the set of *good* segments. If we perform a linear scan on the domain, from begin to end position of the query sequence, then value of S can change only at the 2*n* endpoints of the segments. Therefore, the highest scoring segments overlapping at the 2n endpoints constitute the set of *good* segments, which is precisely what Algorithm 1 computes.

**Algorithm 1:**
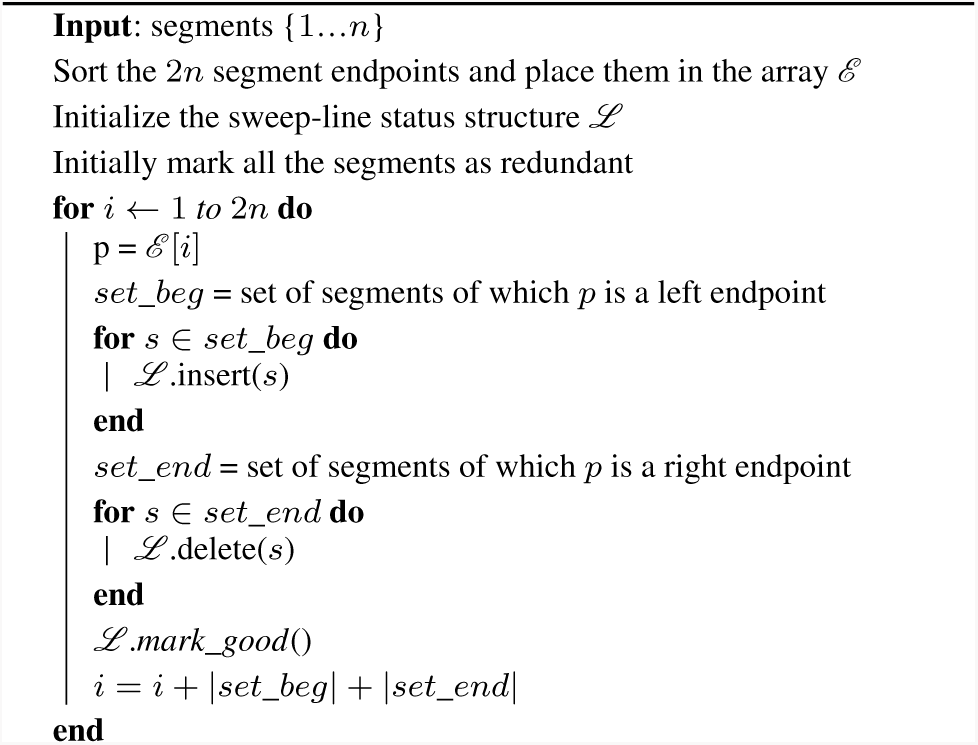
Plane-sweep based alignment filtering algorithm

We make a minor modification to the above described algorithm for efficiency, specifically in the *mark_good* function. In this function, we mark the highest scoring segments in the tree L as *good.* We execute this by traversing the segments in decreasing order in L, starting from the maximum. However, we terminate the traversal if a segment is observed as marked *good* already. This helps avoid redundant computations, and the algorithm still remains correct due to the following property:

##### Lemma 2

Consider all the segments with equal and highest scores in 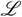: *si, S2,…,Sj*,…, *sk*, ordered in non-increasing manner. Suppose segment s_j_ has been marked good in one of the previous iterations of the algorithm, then the segments *s_j_+i, s_j_+2,…* sk must have already been marked good as well.

Proof. The aforementioned property is satisfied by default during the first iteration of the algorithm because there cannot be any previously marked segments. Suppose this property remains true till iteration i, and we are currently executing iteration i +1. Segments *s_i_, s_2_,… sk € L*, so we know that the sweep line intersects these segments. Also, the ordering of the segments is maintained based on their scores and begin positions, and since the scores of these segments are equal, therefore *begin_pos(s_i_*) > *begin_pos(s_2_) ≥*… ≥ *begin_pos(sk*). Now consider the iteration when segment *sj* was marked *good*. Then, the sweep line must have intersected the segments *sj+_i_,sj+_2_,…sk* as well. Therefore, if the segment *sj* was marked, then the segments s_j_+_i_, *sj+_2_,…sk* must have been marked within or before the same iteration.

The total cost of sorting, *insert* and *delete* operations in Algorithm 1 is clearly *O(n* log *n*). Because the revised *mark_good* function marks at most *n* segments throughout the algorithm, its runtime is also bounded by *O(n* log *n*). Thus, we conclude that the runtime complexity of our alignment filtering algorithm is bounded by *O(n* log *n)* which is restated as a theorem below.

##### Theorem 1

Given n alignment segments, Algorithm 1 solves the alignment filtering problem in *O*(*n log n*) time.

##### Theorem 2

The above proposed filtering algorithm is optimal.

Proof. The INTEGER ELEMENT UNIQUENESS problem (given *n* integers, decide whether they are all unique) is known to have a lower bound of Ω(*n* log *n*) assuming the algebraic decision-tree model (Lubiw and Rácz, 1991). A simple transformation can be designed to show that

INTEGER ELEMENT UNIQUENESS *∝_N_* ALIGNMENT FILTERING

Let *{x_i_, x_2_,…,x_n_}* be a set of *n* integer elements. For each element *xi*, construct a segment with begin position, end position, and score as x_i_, *xi*, and *i* respectively. Because each segment is assigned a unique score, all the *n* elements are unique if and only if the filtering algorithm reports all the segments as *good*.

### 2.3 Related Work for Filtering Alignments

There can be many alternative formulations of the filtering criteria. For instance, BLAST (Altschul *et al.*, 1997) filters out alignments if they are fully contained in ≥ *K* alignments of higher scores (Berman *et al.*, 1999). Berman *et al.* also discussed a weaker alternative filtering condition where a match is filtered out if each position in a segment is covered by ≥ *K* segments of higher score. Note that our filtering formulation is its special case with *K* =1. They discussed a different *O(n* log *n*) algorithm to solve the problem based on interval-tree of all input segments. Although a direct performance comparison is not possible due to unavailability of their implementation, note that the tree size in our plane-sweep based algorithm is limited by the number of overlapping segments which intersect the vertical sweep-line, which can be (and typically is) orders of magnitude smaller than the total count for large datasets. As such, even with the same theoretical complexity, we expect our algorithm to perform faster with less memory usage in practice.

### 2.4 Execution for Mapping Applications

The above filtering criteria is useful to identify the promising alignments between query and reference genomes. For the genome-to-genome mapping application, we execute the filtering algorithm twice, once to filter best alignments for query sequence, followed by filtering best alignments for reference sequence. Mappings which pass both filters constitute the orthologous matches, required for building a one-to-one homology map. For read mapping however, filtering on just the query sequence is appropriate. Accordingly, Mashmap2 provides two filtering modes: one-to-one and map for the two applications respectively.

## 3 Results

We assess the performance of Mashmap2 for genome-to-genome and split-read mapping in comparison to recent versions of state-of-the-art software Minimap2 (Li, 2017) and Nucmer (Marçais G *et al.*, 2018). Results indicate that Mashmap2 provides output of comparable quality, and yields significant gains in memory-usage. Subsequently, we demonstrate the utility of Mashmap2 in accurately computing all 1 Kbp long duplications in the human genome.

### 3.1 Genome-to-Genome Mapping

#### 3.1.1 Datasets

To evaluate and compare Mashmap2 for mapping genome assemblies, we used four datasets D1-D4 listed in Table 1. Dataset D1 includes comparison between microbial genomes E. *coli* O157:H7 and E. *coli* K12. The two instances D2 and D3 require mapping of NA12878 human reference genome assemblies to the hg38 human reference genome. Query genome assemblies in both instances D2 and D3 are the recently published assemblies by Canu (Koren *et al.*, 2017) using ultra-long Oxford Nanopore Technology (ONT) reads (Jain *et al.*, 2017c). Dataset D3 includes a long-read only Canu assembly whereas assembly in dataset D2 is also error-corrected using Illumina reads. Additionally, to evaluate Mashmap2 for the split-read mapping task, D4 includes ultra-long human ONT reads, generated using a single flowcell (Jain *et al.*, 2017c).

**Table 1.**
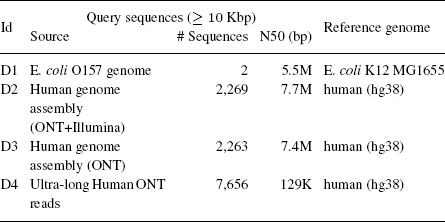
List of datasets used for evaluation. Datasets D1-D3 are included to evaluate Mashmap2 for genome-to-genome mapping application, and D4 for long read mapping application. We discarded a small fraction of contigs and reads which were shorter than 10 Kbp.

#### 3.1.2 Defining Baseline and Methodology

For the purpose of evaluating output accuracy, we used MUMmer package (v4.0.0.beta2), which includes the Nucmer4 alignment program for comparing DNA sequences (Marçais G *et al.*, 2018). Nucmer4 is sensitive enough to report alignments for both assembly and read mapping tasks, therefore we considered its output as truth while evaluating accuracy. Note that computing truth using an exact alignment algorithm is not feasible for the datasets used. We also used Minimap2 (v2.7-r659) (Li, 2017) as a baseline for various performance metrics. Minimap2 executes chaining algorithm on fixed-length exact matches to compute alignment boundaries. To our knowledge, it is among the fastest tools available to map DNA sequences in an alignment-free fashion.

Each software, including ours exposes many parameters (e.g., *k*-mer or seed length). Default *k*-mer size in Mashmap2 is 16. We mostly conform to default parameters with all software tested, except as noted below. Mashmap2 mainly requires a minimum length and identity for the desired local alignments. In this analysis, we targeted long alignments, and accordingly fixed the minimum alignment length requirement as 10 Kbp. We set the minimum alignment identity requirement for the four datasets based on their input characteristics as {D1: 95%, D2: 95%, D3: 90%, D4: 85%}. Accordingly, we tested Mashmap2 for reporting the alignment boundaries as per the provided requirements. Filtering modes were set to one-to-one and map for datasets D1-D3 and D4 respectively. Nucmer4 was run with default parameters, followed by running *delta-filter*, both components of the MUMmer package. Following its user documentation, *delta-filter* was executed with -1 parameter to construct one-to-one alignment map in datasets D1-D3 and -q parameter for read mapping in D4. Finally, Minimap2 supports genome-to-genome mapping mode using -x asm5 flag, and nanopore read mapping mode using -x map-ont. We executed all three software in multi-threaded mode using 8 CPU threads. All comparisons were done on an Intel Xeon E5-2680 platform with 28 physical cores and 256 GB RAM.

#### 3.1.3 Runtime and Memory Usage

The wall-clock runtime and memory-usage of Mashmap2, Minimap2 and Nucmer4 using datasets D1-D4 are shown in Table 2. The runtimes represent end-to-end time, from reading input sequences to generating the final output. Minimap2 can report base-to-base alignments but does not by default. Thus, the final output of Mashmap2 and Minimap2 are alignment boundaries and scores, whereas Nucmer4 outputs base-to-base alignments. We observe that Mashmap2 uses significantly less memory while being competitive in runtime when compared to Minimap2. It improves memory-usage by 5.3x, 4.9x, 4.4x, 2.1x for the four datasets respectively. The performance gap against Nucmer4 is much wider with speedups of 10.4x, 210x, 19.8x, 11.9x, and memory-usage improvements by 8.6x, 15.1x, 14.7x, 10.6x on the datasets D1-D4 respectively.

**Table 2.** Total execution time and memory usage comparison of Mashmap2 against Minimap2 and alignment-based tool Nucmer4. All software were run in parallel using 8 CPU threads.

| Id | Mashmap2 |  | Minimap2 |  | Nucmer4 |  |
| --- | --- | --- | --- | --- | --- | --- |
|  | Time | Memory | Time | Memory | Time | Memory |
| D1 | 0.5s | **16M** | **0.4s** | 85M | 5.2s | 138M |
| D2 | **1m 26s** | **3.5G** | 3m 3s | 17.3G | 5h 1m | 53G |
| D3 | 6m 33s | **3.6G** | **3m 11s** | 15.9G | 2h 10m | 53G |
| D4 | **2m 6s** | **5.0G** | 3m 10s | 10.3G | 25m 2s | 53G |

Mashmap2 and Minimap2 follow the same initial step of sampling *k*-mers using minimizers (Schleimer *et al.*, 2003; Roberts *et al.*, 2004), followed by computing their exact matches in the reference genome.

However, they differ in using these exact matches to compute the mappings. Mashmap2 includes an efficient MinHash-based mechanism to estimate Jaccard similarity and auto-tunes the internal parameters (e.g., *k*-mer sampling rate, Jaccard similarity threshold), conforming to the local alignment identity and length requirements provided by user. Auto-tuning can help reduce memory-usage and achieve faster runtime with increasing identity and length thresholds. For instance, Mashmap2 achieves better performance on dataset D2 than D3 because the error-rate of input sequences in D2 is lower due to the error-correction using Illumina reads. Based on results of Jain et *al.* (2017c), the error-correction phase improved average alignment identity from 95% to 99.3% with respect to the hg38 reference. It is important to maintain high accuracy while being fast, therefore we next evaluate the quality of output.

#### 3.1.4 Accuracy

Accuracy evaluation of Mashmap2 and Minimap2 in comparison with alignments produced by Nucmer4 is shown in Table 3. Recall was measured against Nucmer4 output alignments which satisfy the alignment requirements in terms of minimum length and identity provided to Mashmap2. We also expected Minimap2 to report these alignments because it is designed to compute matches in these identity ranges.

A reported local alignment boundary estimate by Mashmap2 or Minimap2 was assumed to recall a Nucmer4 alignment if it overlapped with the alignment on both query and reference sequences, and if the mapping strand matched. From Table 3, we observe that both Mashmap2 and Minimap2 consistently achieved high recall scores, with Minimap2 performing slightly better. Obtaining high recall scores by itself is not sufficient, because it can be achieved by mapping a query sequence to all possible positions. In parallel to achieving high recall scores, both Mashmap2 and Minimap2 mapped a large fraction of query genome assemblies to unique mapping positions in the reference genomes. To show this, we computed the fraction of base-pairs of the query sequence that are mapped to a single position on the reference genome (Table 3). Finally, Mashmap2 provides a script to visualize its output as dot-plots, similar to MUMmerplot. These dot-plots when visually inspected, appeared similar to Nucmer4's output. Here we show homology maps computed by Mashmap2 using datasets D1 and D2 (Figure 4).

**Table 3.** Accuracy evaluation of Mashmap2 and Minimap2 to do an alignment-free computation of mapping boundaries. Recall was measured while assuming Nucmer4 output alignments as truth.

| Id | Recall scores |  |  | Fraction of query bases mapped uniquely |  |
| --- | --- | --- | --- | --- | --- |
|  | Mashmap2 | Minimap2 | Count of Nucmer4 alignments | Mashmap2 | Minimap2 |
| D1 | **100%** | **100%** | 144 | 74.0% | **78.9%** |
| D2 | 97.5% | **98.3%** | 35,186 | **96.8%** | 96.3% |
| D3 | 97.1% | **98.1%** | 37,807 | **96.9%** | 96.0% |
| D4 | 99.3% | **99.5%** | 4,349 | 81.2% | **84.6%** |

**Fig. 4.**
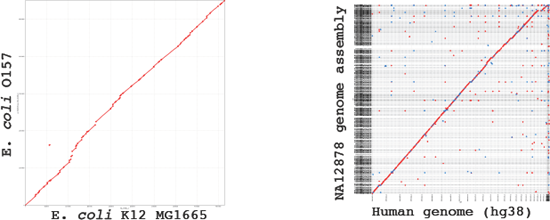
Dot plots for genome to genome mappings for datasets D1 and D2 generated using Mashmap2. Mashmap2 supports visualization using MUMmerplot tool, modified to be compatible with its output format.

#### 3.1.5 Efficacy of The Filtering Algorithm

Eukaryotic genomes tend to contain a lot of repetitive sequences, therefore, the motivation behind our plane-sweep based filtering heuristic is to discard noisy mappings, and compute promising matches between the query and reference genomes. We show the importance and effectiveness of our filtering strategy in Table 4. Note that a large fraction of mappings was pruned out by the filter. While doing so, high recall scores against Nucmer4 alignments were maintained (see Table 4). Although we do not present the contribution of this phase to the total runtime, our plane-sweep algorithm is fast in practice; it used an insignificant fraction of the total runtime.

**Table 4.** Effectiveness of the filtering algorithm in Mashmap2. A large fraction of mappings were filtered out by the algorithm, while the recall scores against the Nucmer4 alignments remained largely unaffected. Last column in this table is copied from Table 3 for convenience.

| Id | Count of output mappings |  |  | Recall scores |  |
| --- | --- | --- | --- | --- | --- |
|  | Without filter | With filter | Ratio (without/with) | Without filter | With filter |
| D1 | 145 | 82 | 1.77 | 100.0% | 100.0% |
| D2 | 6,541,930 | 3,985 | 1,642 | 99.9% | 97.5% |
| D3 | 53,331,538 | 3,137 | 17,001 | 99.7% | 97.1% |
| D4 | 3,881,667 | 15,311 | 254 | 99.9% | 99.3% |

### 3.2 Computing Duplications in the Human Genome

Soon after the publication of the human genome, it was realized that the genome is replete with repetitive sequences (International Human Genome Sequencing Consortium, 2004). Intra- and inter-chromosomal duplications have been found to play a vital role in genome evolution, its stability, and diseases (Emanuel and Shaikh, 2001), and knowing the location of such repeats can be important for many genomic analyses. Yet, fully annotating all repeats in a genome can be computationally intensive. To demonstrate the scalability of Mashmap2, we computed all ≥ 1 Kbp duplications in the human genome (GRCh38, Schneider et al., 2017) with ≥ 90% alignment identity. Due to probabilistic guarantees, we expect Mashmap2 to estimate the boundaries of all such duplications with a high recall value. Typical genome-to-genome aligners do not provide such guarantees, and may require extensive parameter tuning. The importance of these duplications has been known for a long time (Emanuel and Shaikh, 2001; *Bailey et al., 2002*); accordingly the UCSC genome browser (*Kent et al., 2002*) also maintains them in a database (named as segmental duplications) for the human genome. We summarize our method below and contrast our output with this database.

#### 3.2.1 Methodology

We used 24 chromosome sequences (1-22, X,Y) and mitochondrial DNA from the hg38 version of the human genome as our input sequence set. To compute all ≥ 1 Kbp, ≥90% identity duplications, we executed Mashmap2 with the same length and identity requirements, with filtering disabled. Mashmap2 reported 8.4 billion alignment boundaries for all duplications after finishing its run. The count of reported mappings is high due to several high-copy repeat families in the genome, not all of which exceed our minimum thresholds. To remove the shorter or lower identity mappings, each of the approximate alignments was processed using LAST (Kiełbasa *et al.*, 2011) to compute a base-level alignment. This resulted in 213.9 million validated alignments ≥ 1 Kbp in length and ≥ 90% identity. Finally, we filtered out trivial duplications (i.e. regions matching with themselves), and were left with 213.8 million alignments. This experiment took 120 CPU hours for executing Mashmap2 and 175,000 CPU hours for validating all reported mappings using LAST. We converted these alignments into BED format to compare against the UCSC database using Bedtools (Quinlan and Hall, 2010); the results are discussed below.

#### 3.2.2 Accuracy Evaluation and Insights

The UCSC Segmental Duplications database for the hg38 human genome was computed using a method proposed by Bailey et *al.* (2001), and was last updated in 2014. It is important to note that prior to computing genomic duplications, their method removed high-copy repeat elements (e.g., LINEs, *Alus)* from the genome. Therefore, this database does not constitute all ≥1 Kbp, ≥90% identity duplications in the genome, but a significant fraction of them. Nevertheless, we should expect our output to have high recall against this database. To measure recall on each chromosome, we computed coverage of those UCSC duplication annotations that have overlap with Mashmap2 duplications, and divided it by the coverage of all UCSC duplication annotations. Therefore, a 100% recall score would imply that all base-pairs which are annotated as segmental duplication in the UCSC database are part of one or more Mashmap2 alignments. We show these recall scores for each chromosome as well for the complete genome in Figure 5. Recall is consistently observed to be above 90% for each chromosome, and the aggregate recall for the complete genome is 97.2%. High recall scores achieved here, as well as in our prior experiments, demonstrate high sensitivity of our algorithm in practice.

**Fig. 5.**
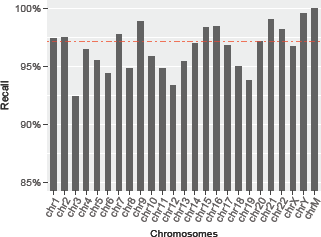
Recall scores of duplications computed using Mashmap2 against the UCSC segmental duplication database. Above 90% recall scores are achieved on each chromosome consistently. The red dotted line shows the aggregate recall score of 97.2% for the complete genome.

Finally, we compared the coverage of our alignments versus the UCSC database. Since our method did an exhaustive search of all duplications with ≥ 1 Kbp length and ≥ 90% identity without masking any genomic repeats, we observe that our algorithm attains either equal or higher coverage on each chromosome (Figure 6). For the complete genome, coverage of our alignments is 10.4%; 5% higher than the coverage of UCSC annotations. We examined the subset of our duplications which do not overlap with UCSC segmental duplications. Indeed a large coverage fraction (81%) comprises of high-copy repeats (i.e, coverage depth ≥ 50), potentially due to common repeat elements. However, a significant coverage (1.03% of complete genome) is composed of low-copy repeats, with coverage depth ≤ 50 indicating the potential to uncover novel segmental duplications. Validating this possibility requires a more careful inspection of the output, and will be our future work.

**Fig. 6.**
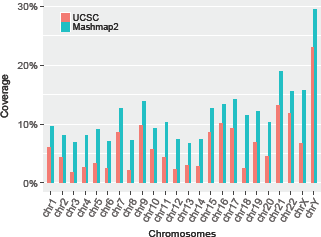
Comparison of genomic coverage between the UCSC Segmental Duplication database and Mashmap2 output alignments. Both methods reported equal coverage 83% on mitochondrial chromosome (not shown above to keep the plot legible). Coverage of duplications computed using our method is significantly higher, owing to its exhaustive search of all repeats with ≥ 1 Kbp length and ≥90% identity without repeat masking.

## 4 Discussion

In this work, we presented a fast algorithm for computing homology maps between whole genomes. We have given both theoretical and experimental evidence of the sensitivity provided, in terms of computing local alignment boundaries based on the minimum alignment length and identity parameters. This formulation grants a convenient mechanism for users to execute this algorithm based on the underlying applications, which can be (but not limited to) mapping genome assembly of variable quality, aligning long reads to reference genomes, or computing segmental duplications in large genomes. Additionally, we formulated a filtering heuristic, and proposed an optimal plane-sweep based filtering algorithm for prioritizing alignments based on their scores and locations. The filtering algorithm is practically fast, accurate, and easy to implement in few lines of code using standard libraries. When mapping a human genome assembly to the human reference genome, Mashmap2 takes only about a minute from reading input sequences to generating the final alignment boundaries, identity estimates, and a dot-plot for visualization. Because of the underlying auto-tuning mechanism in Mashmap2, performance depends on the sensitivity requirements provided to the algorithm. As the pace of whole-genome sequencing continues to increase, faster practical algorithms and theoretical advances will help analyze available and forthcoming data.

Although our algorithm accelerates mapping of a single genome assembly to a single reference genome, its runtime would scale linearly when mapping to multiple reference genomes. Future work will include development of sub-linear algorithms using existing ideas of non-linear reference genome representations. We also plan to evaluate biological novelty of the human segmental duplications computed in this work.

## Acknowledgements

We thank Pavel Pevzner for motivating evaluation of segmental duplications. This research was supported in part by the Intramural Research Program of the National Human Genome Research Institute, National Institutes of Health, and the U.S. National Science Foundation under IIS-1416259. We also acknowledge the use of computing resources provided through the Partnership for an Advanced Computing Environment (PACE) at the Georgia Institute of Technology, and the Biowulf system at the National Institutes of Health, Bethesda, MD (https://hpc.nih.gov/).

## References

Altschul, S. F., Madden, T. L., Schäffer, A. A., Zhang, J., Zhang, Z., Miller, W., and Lipman, D. J. (1997). Gapped BLAST and PSI-BLAST: a new generation of protein database search programs. Nucleic acids research, 25(17), 3389–3402.

Bailey, J. A., Yavor, A. M., Massa, H. F., Trask, B. J., and Eichler, E. E. (2001). Segmental duplications: organization and impact within the current human genome project assembly. Genome research, 11(6), 1005–1017.

Bailey, J. A., Gu, Z., Clark, R. A., Reinert, K., Samonte, R. V., Schwartz, S., Adams, M. D., Myers, E. W., Li, P. W., and Eichler, E. E. (2002). Recent segmental duplications in the human genome. Science, 297(5583), 1003–1007.

Berman, P., Zhang, Z., Wolf, Y. I., Koonin, E. V., and Miller, W. (1999). Winnowing sequences from a database search. In Proceedings of the third annual international conference on Computational molecular biology, pages 50–58. ACM.

Bray, N., Dubchak, I., and Pachter, L. (2003). AVID: A global alignment program. Genome research, 13(1), 97–102.

Brudno, M., Chapman, M., Göttgens, B., Batzoglou, S., and Morgenstern, B. (2003). Fast and sensitive multiple alignment of large genomic sequences. BMC bioinformatics, 4(1), 66.

Delcher, A. L., Kasif, S., Fleischmann, R. D., Peterson, J., White, O., and Salzberg, S. L. (1999). Alignment of whole genomes. Nucleic acids research, 27(11), 2369–2376.

Emanuel, B. S. and Shaikh, T. H. (2001). Segmental duplications: an’expanding’role in genomic instability and disease. Nature Reviews Genetics, 2(10), 791–800.

Grabherr, M. G., Russell, P., Meyer, M., Mauceli, E., Alföldi, J., Di Palma, F., and Lindblad-Toh, K. (2010). Genome-wide synteny through highly sensitive sequence alignment: Satsuma. Bioinformatics, 26(9), 1145–1151.

Haussler, D., O’Brien, S. J., Ryder, O. A., Barker, F. K., Clamp, M., Crawford, A. J., Hanner, R., Hanotte, O., McGuire, J. A., Miller, W., et al. (2009). Genome 10K: a proposal to obtain whole-genome sequence for 10 000 vertebrate species.

International Human Genome Sequencing Consortium (2004). Finishing the euchromatic sequence of the human genome. Nature, 431(7011), 931–945.

Jain, C., Dilthey, A., Koren, S., Aluru, S., and Phillippy, A. M. (2017a). A fast approximate algorithm for mapping long reads to large reference databases. In International Conference on Research in Computational Molecular Biology, pages 66–81. Springer.

Jain, C., Rodriguez-R, L. M., Phillippy, A. M., Konstantinidis, K. T., and Aluru, S. (2017b). High-throughput ANI analysis of 90K prokaryotic genomes reveals clear species boundaries. bioRxiv, page 225342.

Jain, M., Koren, S., Quick, J., Rand, A. C., Sasani, T. A., Tyson, J. R., Beggs, A. D., Dilthey, A. T., Fiddes, I. T., Malla, S., et al. (2017c). Nanopore sequencing and assembly of a human genome with ultra-long reads. bioRxiv, page 128835.

Kent, W. J., Sugnet, C. W., Furey, T. S., Roskin, K. M., Pringle, T. H., Zahler, A. M., and Haussler, D. (2002). The human genome browser at UCSC. Genome research, 12(6), 996–1006.

Kiełbasa, S. M., Wan, R., Sato, K., Horton, P., and Frith, M. C. (2011). Adaptive seeds tame genomic sequence comparison. Genome research, 21(3), 487–493.

Koren, S., Walenz, B. P., Berlin, K., Miller, J. R., Bergman, N. H., and Phillippy, A. M. (2017). Canu: scalable and accurate long-read assembly via adaptive k-mer weighting and repeat separation. Genome research, 27(5), 722–736.

Kurtz, S., Phillippy, A., Delcher, A. L., Smoot, M., Shumway, M., Antonescu, C., and Salzberg, S. L. (2004). Versatile and open software for comparing large genomes. Genome biology, 5(2), R12.

Lander, E. S., Linton, L. M., Birren, B., Nusbaum, C., Zody, M. C., Baldwin, J., Devon, K., Dewar, K., Doyle, M., FitzHugh, W., et al. (2001). Initial sequencing and analysis of the human genome. Nature, 409(6822), 860–921.

Li, H. (2017). Minimap2: fast pairwise alignment for long DNA sequences. arXiv preprint arXiv:1708.01492.

Lubiw, A. and Rácz, A. (1991). A lower bound for the integer element distinctness problem. Information and Computation, 94(1), 83–92.

Ma, B., Tromp, J., and Li, M. (2002). Patternhunter: faster and more sensitive homology search. Bioinformatics, 18(3), 440–445.

Marçais G, Delcher, A. L., Phillippy, A. M., Rachel, C., Salzberg, S. L., and Aleksey, Z. (2018). MUMmer4: A fast and versatile genome alignment system. PLoS Comput Biol, 14(1).

Ondov, B. D., Treangen, T. J., Melsted, P., Mallonee, A. B., Bergman, N. H., Koren, S., and Phillippy, A. M. (2016). Mash: fast genome and metagenome distance estimation using MinHash. Genome biology, 17(1), 132.

Quinlan, A. R. and Hall, I. M. (2010). Bedtools: a flexible suite of utilities for comparing genomic features. Bioinformatics, 26(6), 841–842.

Roberts, M., Hayes, W., Hunt, B.R., Mount, S. M., and Yorke, J. A. (2004). Reducing storage requirements for biological sequence comparison. Bioinformatics, 20(18), 3363–3369.

Schleimer, S., Wilkerson, D. S., and Aiken, A. (2003). Winnowing: local algorithms for document fingerprinting. In Proceedings of the 2003 ACM SIGMOD international conference on Management of data, pages 76–85. ACM.

Schneider, V. A., Graves-Lindsay, T., Howe, K., Bouk, N., Chen, H.-C., Kitts, P. A., Murphy, T. D., Pruitt, K. D., Thibaud-Nissen, F., Albracht, D., et al. (2017). Evaluation of grch38 and de novo haploid genome assemblies demonstrates the enduring quality of the reference assembly. Genome research, 27(5), 849–864.

Schwartz, S., Kent, W. J., Smit, A., Zhang, Z., Baertsch, R., Hardison, R. C., Haussler, D., and Miller, W. (2003). Human-mouse alignments with BLASTZ. Genome research, 13(1), 103–107.

Shamos, M. I. and Hoey, D. (1976). Geometric intersection problems. In Foundations of Computer Science, 1976., 17th Annual Symposium on, pages 208–215. IEEE.

Venter, J. C., Adams, M. D., Myers, E. W., Li, P. W., Mural, R. J., Sutton, G. G., Smith, H. O., Yandell, M., Evans, C. A., Holt, R. A., et al. (2001). The sequence of the human genome. science, 291(5507), 1304–1351.

Vyverman, M., De Baets, B., Fack, V., and Dawyndt, P. (2013). essamem: finding maximal exact matches using enhanced sparse suffix arrays. Bioinformatics, 29(6), 802–804.

